# Liver-derived cell lines from cavefish *Astyanax mexicanus* as an in vitro model for studying metabolic adaptation

**DOI:** 10.1101/2022.01.06.475101

**Authors:** Jaya Krishnan, Yan Wang, Olga Kenzior, Hassan Huzaifa, Luke Olsen, Dai Tsuchiya, Yongfu Wang, Chongbei Zhao, Nicolas Rohner

## Abstract

Cell lines have become an integral resource and tool for conducting biological experiments ever since the Hela cell line was first developed (1). They not only allow detailed investigation of molecular pathways but are faster and more cost-effective than most in vivo approaches. The last decade saw many emerging model systems strengthening basic science research. However, lack of genetic and molecular tools in these newer systems pose many obstacles. *Astyanax mexicanus* is proving to be an interesting new model system for understanding metabolic adaptation. To further enhance the utility of this system, we developed liver-derived cell lines from both surface-dwelling and cave-dwelling morphotypes. In this study, we provide detailed methodology of the derivation process along with a comprehensive biochemical and molecular characterization of the cell lines, which reflects key metabolic traits of cavefish adaptation. We anticipate these cell lines to become a useful resource for the *Astyanax* community as well as researchers investigating fish biology, comparative physiology, and metabolism.

## Introduction

*Astyanax mexicanus* has emerged as a powerful system to understand the genetic and physiological basis of metabolic adaptation to nutrient deprived habitats (2–10). The species comes in two morphotypes – the surface fish that live in lush, nutrient-rich rivers and the blind cavefish that dwell in dark and nutrient-poor caves in the Sierra de El Abra of northeast Mexico. The cavefish survive the nutrient-deprived conditions by remodeling their metabolism accumulating fat stores and becoming hyperglycemic when food is available (8,10). The liver plays an essential role in this process as a key organ for integration of fat and glucose metabolism and thereby, metabolic adaptation. Previous studies from our lab have shown that cavefish have fatty livers and show differential regulation of many key metabolic genes combined with massive differences in chromatin architecture that correlate with gene expression changes (4,8). Liver cell cultures from several organisms are now commercially available including multiple cell lines from human, mouse, zebrafish, rainbow trout, etc. These in vitro systems have proven invaluable for many biochemical, toxicology and cell biology experiments. With the availability of gene editing technologies and high-throughput reporter assays enabling rapid and detailed characterization of genes and regulatory elements, there is a need to develop cell lines for *Astyanax mexicanus.* To this end, we have generated cell lines from surface fish livers (SFL) and Pachón cavefish livers (CFL).

The SFL and CFL are adherent cells, have epithelial cell-like morphology and can be easily grown and maintained in cell culture facility. In this study, we have characterized the cell lines for multiple parameters such as liver enzyme activity, karyotyping and transcriptome. The cells simulate several features that are seen in the fish liver physiology and thus are set to become a useful in vitro model to study the genetic and molecular underpinnings of metabolic adaptation of *Astyanax mexicanus.*

## Methods

### *Astyanax* husbandry

*Astyanax* are housed in polycarbonate or glass fish tanks on racks (Pentair, Apopka, FL) with a 14:10 h light:dark photoperiod. Each rack uses an independent recirculating aquaculture system with mechanical, chemical, and biologic filtration, and UV disinfection. Water quality parameters are maintained within safe limits (upper limit of total ammonia nitrogen range 1 mg/L; upper limit of nitrite range 0.5 mg/L; upper limit of nitrate range 60 mg/L; temperature set-point of 22°C; pH 7.65, specific conductance 800 μS/cm; dissolved oxygen >90%. Water changes range from 20-30 % daily (supplemented with Instant Ocean Sea Salt [Blacksburg, VA]). Adult fish are fed three times a day during breeding weeks and once per day during non-breeding weeks on a diet of Mysis shrimp (Hikari Sales USA, Inc., Hayward, CA) and Gemma 800 (Skretting USA, Tooele, UT).

### Media composition

The complete media composition was adopted from the media used for a zebrafish liver cell line with minor changes (11), consisting of 45% Leibovitz’s L-15, 30% Dulbecco’s modified Eagle’s and 15% Ham’s F12, 15mM Hepes-NaOH, 0.01mg/ml bovine insulin, 50ng/ml mouse epidermal growth factor, 5% heat-inactivated fetal bovine serum, 0.5% Trout serum and 1% embryo extract. The embryo extract was used for the initial ~10-15 passages until the cell lines stabilized. Extracts from 3dpf surface fish was used for SFL and 3dpf Pachón fish for CFL. Embryo extract was made as follows: 3dpf embryos were collected (minimum 200 embryos) and rinsed with Ringer’s solution. The embryos were then homogenized in Leibovitz’s L-15 media supplemented with 2x Penicillin-Streptomycin (PS) (Thermo Fisher #10378016) (1ml media per 200 embryos) using the tight pestle of a Dounce homogenizer. The extract was centrifuged to remove the debris and the supernatant collected and frozen in 1 ml aliquots at −20°C.

### Liver tissue preparation and primary cell derivation

The cell line derivation was done from livers of ~150 days old fish. While the surface fish were not pre-conditioned before dissection, we starved Pachón fish for one week prior to liver dissection. As cavefish livers store more fat than the surface fish livers, we believe the starvation reduced the fat content in the livers which helped in the overall survival of the cells in vitro. The fish were euthanized in MS-222 and livers were dissected. The tissues, each weighing around 20mg, were then quickly rinsed once in 70% alcohol and washed multiple times with phosphate buffered saline (PBS) supplemented with 500μg/ml of Penicillin-Streptomycin cocktail (final concentration of 5X). Each well of a 6-well plate contained 1-2 livers. The tissue was minced with a scalpel and trypsinized at room temperature (RT) for 5 min. Tissue was further disintegrated by pipetting up and down about 20 times with a 1ml pipette tip during the incubation. The dissociated cells were then transferred to 15 ml conical tubes. Cells from one well were split to 2×15ml tubes and pelleted at RT at 1000 rpm for 4 min. The cells were resuspended in 1ml of ZFL-c + 1x PS + 5% embryonic extract. 1ml of cell suspension was seeded in one well of 48-well plate (gelled with 0.1% Gelatin at RT for 0.5 hour) (Note: Cells from one liver (~20mg) can be seeded to 2 wells of 48-well plate, approximately).

### Cell line establishment

We replaced half of the culture media in each well with fresh media after 24 hours. The media was changed every 2 days (we replaced only half of the old media to provide 50% conditioned media to the cells). Cells were passaged in a 1:2 ratio every week. Once the cells reached the T-25 flask, 1-2 x 10^6^ cells were frozen for back up (Note: massive cell death is observed initially but a significant number of cells will attach and eventually grow). To obtain homogeneous cell populations, cell cloning by serial dilution method was performed according to the standard protocol from Corning (corning.com/catalog/cls/documents/protocols/Single_cell_cloning_protocol.pdf). In brief, the heterogeneous cells were serially diluted in 96-well plates and single cell colonies were picked once they became large and conspicuous. The SFLs were at passage #37 and the CFLs at #33 when the serial dilutions were performed. Once the clones reached 6-well plates, clones that had similar and homogeneous morphology between SFL and CFL were chosen for further characterization.

### Preparation of metaphase spreads and chromosome counting

The cells were seeded in T-25 flasks to a density of 1e6 the day before chromosome preparation. On the next morning, 0.25% final concentration of colchicine was added to the cells and incubated for 2 hours. The cells were collected and washed in PBS, centrifuged at 300 x g for 3 min at RT. Hypotonic solution (0.075M KCl) was applied to the cell pellet, incubated for 15 min at RT, and centrifuged at 188 x g for 3 min at RT. Freshly prepared ice-cold fixative (75% Methanol, 25% glacial acetic acid) was added to the cell pellet, and the tube was kept on ice for 10 min. The suspension was centrifuged at 188 x g for 5 min at 4°C, and the pellet was resuspended in cold fixative. After 10 min incubation on ice, the suspension was centrifuged, resuspended in cold fixative one more time, and stored at −20°C until use. For the preparation of metaphase spreads, the cell suspension was dropped on a slide, and the slide was dried on a heat block at 75°C for 3 minutes. Slides were stained with DAPI (10μg/ml) for 10min, and Prolong gold was applied as a mounting media. For chromosome counting, at least 88 metaphase spreads were analyzed using Zeiss Axio Vert 200M fluorescence microscope (Carl Zeiss, Jena, Germany) equipped with a 63X N.A. 1.4 oil-immersion objective (Carl Zeiss, Jena, Germany). Image analysis and processing were performed using ImageJ software.

### Seahorse assay

We followed the procedure described in the Seahorse Glycolysis Stress Test kit. In brief, 30,000 cells were seeded in each well of a 96-well plate a day prior to the day of experiment. Each cell line had 8-16 replicates on the same plate. On the day of experiment, the plate was placed in the BioTek Cytation system for brightfield imaging to count cells for normalization of the results. The plate was next transferred to the microplate stage of a Seahorse XFe96 flux analyzer (Seahorse). The initial extracellular acidification rate (ECAR) measurements were taken in the absence of glucose using a 3-minute mix and 3-minute read cycling protocol. Three separate readings were taken to ensure stability. Next, glucose was added to each well to a concentration of 10 mM, and three separate ECAR readings were taken. This was followed by an injection of oligomycin and Hoechst 33342. The final concentration of oligomycin in each well was 2 μM, and three separate ECAR readings were taken. Next, 2-deoxyglucose was injected to a final concentration of 25 mM in each well, and three separate ECAR readings were taken. XFe data normalization was performed by in situ nuclear staining and in situ cell counting using the imaging and normalization system (BioTek Cytation).

### ALT assay

Cells were grown to 80% confluency and 1 x 10^6^ cells were collected as pellets in duplicates for the assay. Colorimetric assay to measure alanine aminotransferase activity in the cell lines was performed using the ALT activity assay kit from Sigma (MAK052-1KT) according to the manufacturer’s instructions. Zebrafish Liver cell line was used as positive control.

### Antibiotic kill curve

We plated cells at 90% per well of a 12-well plate. We kept 1 well in maintenance media as control. We set the remaining wells in chosen selection concentrations and ensured that the cells are from a healthy, routinely sub-cultured stock. We observed the culture daily and changed media with corresponding antibiotic concentrations. We determined which concentration to use by observation and choose the concentration 100μg/ml of G418 above the concentration that has 100% cell death.

### RNA-seq

Alignment and differential expression: RNA-seq reads were aligned to *Astyanax mexicanus* reference genome from University of California at Santa Cruz using STAR version 2.7.3 with gene model retrieved from Ensembl, release 102 to generate gene read counts. The transcript abundance ‘TPM’ (Transcript per Million) was quantified using RSEM version 1.3. Differentially expressed genes were determined using R package edgeR version 3.30.3 after filtering low expressed genes with a CPM (Counts Per Million) of 0.2 in at least one library. The resulting p-values were adjusted with Benjamini-Hochberg method using R function p.adjust. Genes with an adjusted p-value < 0.05 and a fold change of 2 were termed as differentially expressed.

Functional enrichment or gene ontology (GO) analysis: Gene functional enrichment analysis was performed on the differentially expressed genes identified in edgeR. A custom wrapper around R package ‘clusterProfiler’ with gene-GO terms retrieved from Ensembl BioMart was used to identify overrepresented GO terms in the differentially expressed genes compared with the background list of all genes.

## Results

We derived cell lines from surface fish and Pachón cavefish livers using dissected livers from 5-month-old fish. The initial set of derived cell population was very heterogeneous, and we performed serial dilutions to isolate clonal cell populations. From the primary derivation, the cell lines have now been in culture for >60 passages. The clonal populations were derived at passage #37 for SFL and #33 for CFL. The cells have an epithelial cell-like morphology and adhere to the culture plate surface (Figure 1 a, b). They are easy to maintain in a media containing DMEM, L-15, Ham’s F-12, trout, and fetal bovine serums, supplemented with insulin and epidermal growth factor and are passaged every week. Initial passages required supplementing the media with 5% *Astyanax* embryo extract (see methods for details), however, once cultures became stable (passage 17 for SFL and passage 12 for CFL) the embryo extract was removed without affecting cell survival. All other components were found to be indispensable for cell survival. For routine passaging, the cells were trypsinized with TrypLE express (Thermo Fischer) for 3-4 min and pelleted by centrifugation at 130g, washed with PBS and seeded in desired density. We observed the growth curve for the two cell lines and calculated their doubling times using standard formula (Roth V. 2006 Doubling Time Computing, Available from: http://www.doubling-time.com/compute.php). Both the cell lines had a comparable doubling time with SFL dividing ~13% faster than CFL on average.

**Figure 1:**
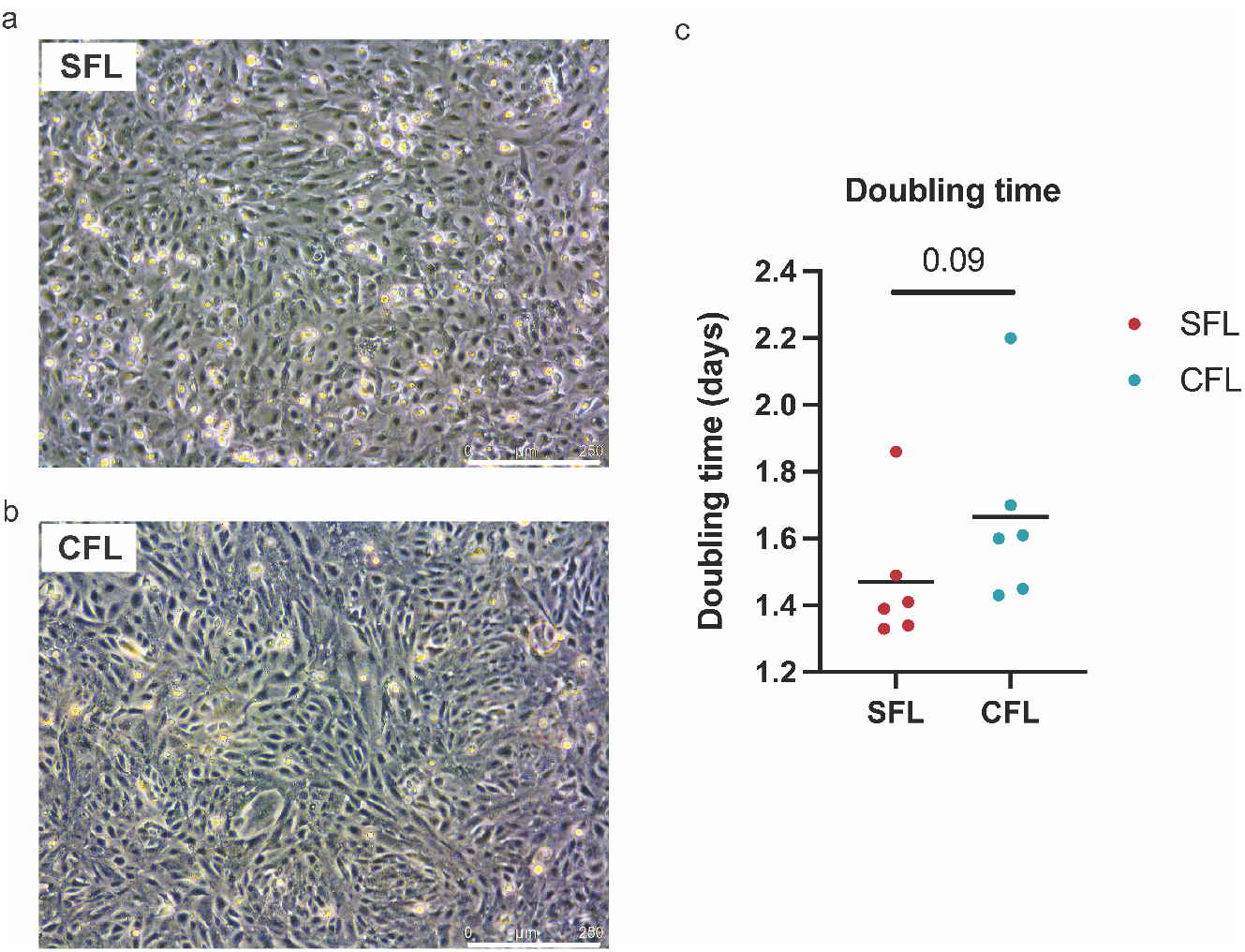
Brightfield images of SFL (a) and CFL (b). The scale denotes 250μM. (c) Graph showing the calculated doubling time for the two cell lines.

### Karyotyping

Metaphase chromosome spreads were made for karyotyping the cell lines. We observed a modal chromosome number of 50 (haploid chromosome number n = 25) for the SFL which was equal to the physiological chromosome count as revealed by genome sequencing (12). The CFL cell line was hyperdiploid with a modal chromosome count of 58 (haploid chromosome number n=29). A recent study showed that Pachón males have B chromosomes (13) and we suspect that some of these extra chromosomes could be B chromosomes.

### SFL and CFL express liver-specific enzymes

Liver tissue is characterized by the production of liver-specific enzymes such as alanine aminotransferase (ALT) and aspartate aminotransferase. To test for liver-specific activity levels in these cell lines, we performed ALT assay and used a zebrafish liver cell line (ZFL) as control (See Supplementary Figure 1 for standard curves). We observed ALT activity in both the cell lines (Table 1).

**Table 1:** The table shows the ALT activity levels of the cell lines.

| Cell line | Activity |
| --- | --- |
| SFL | 3.940 milliunits/ml |
| CFL | 1.577 milliunits/ml |
| ZFL | 0.882 milliunits/ml |

### CFL have increased glycolytic capacity

Previous studies have shown drastic metabolic differences in the metabolism of surface fish and cavefish (4,5,14). Most of the differences that have been documented affect glucose and fat metabolism pathways. For example, it has been shown that surface fish have increased fat catabolism while cavefish have increased fat anabolism (4,14). Furthermore, metabolomic analysis of liver tissues show that cavefish have increased glycolysis as compared to surface fish (5). To test if the energy metabolism of the corresponding cell lines shows similar differences between the morphotypes, we performed two rounds of glycolytic stress test using the Seahorse Analyzer according to the manufacturer’s instruction and published protocols (15) (Figure 3). The analyzer measures the Extracellular Acidification Rate (ECAR) which is the rate of decrease in pH in the assay media. It also provides the Oxygen Consumption Rate (OCR) which is the rate of decrease of oxygen concentration in the assay media. OCR gives a measure of the rate of mitochondrial respiration of the cells. Upon addition of oligomycin, a chemical that inhibits aerobic respiration causing the cells to rely on glycolysis for energy, the cavefish cells showed a higher rate of metabolism indicating their ability for increased glycolysis (Figure 3a-d). This is also depicted clearly in the OCR vs ECAR graph (Figure 3e, f). While the glycolytic capacity of the two cell types was not drastically different, we observed increased glycolysis in the cavefish cell line in 2 independent experiments with 8-16 technical replicates. The results underscore the utility of these cell lines for future metabolic characterization.

**Figure 2:**
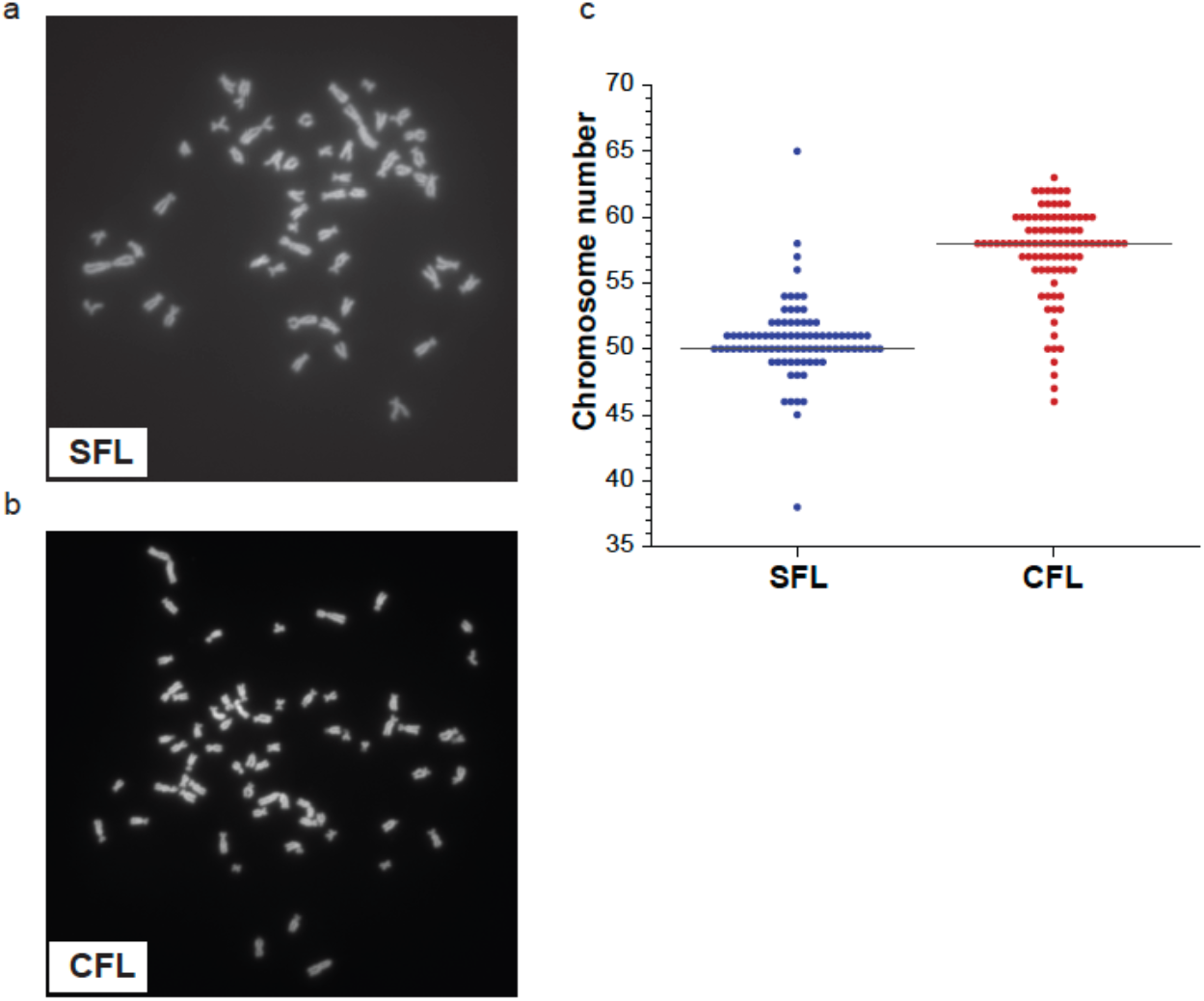
(a) Representative chromosome spread for SFL. (b) Representative chromosome spread for CFL. (c) The graph shows the chromosome counting for 88 and 92 spreads respectively for SFL and CFL.

**Figure 3:**
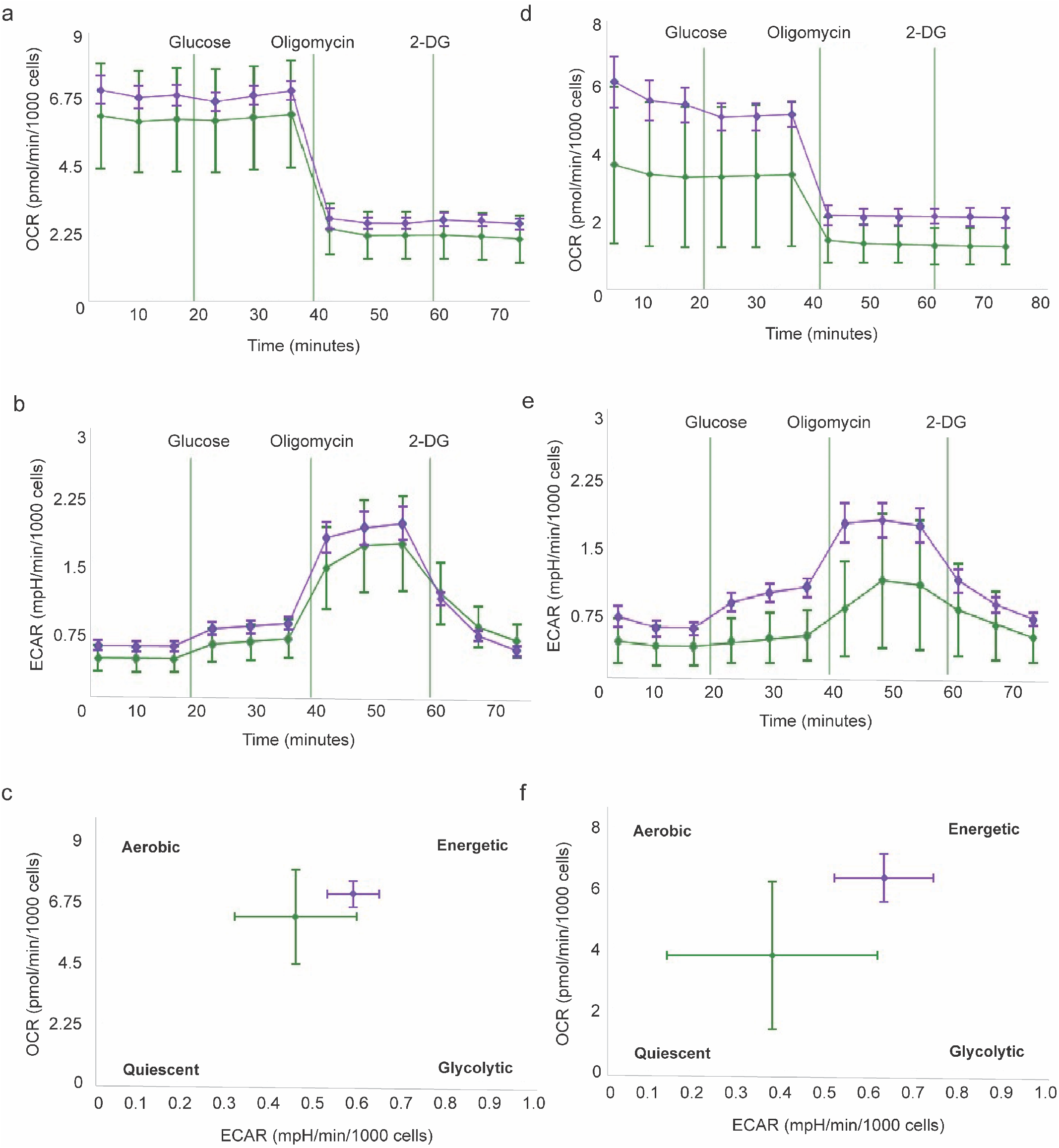
The figure shows results from the Seahorse assay. Left and right panels depict two independent experiments performed using identical procedures. (a, b) Oxygen consumption rate (OCR) for SFL and CFL. (c, d) Extracellular acidification rate (ECAR) for SFL and CFL. (e, f) OCR vs ECAR graph reflecting increased glycolytic capacity for CFL.

### Transcriptome analysis shows increased lipid catabolism in SFL

Next, to characterize the gene expression in the cell lines, we analyzed their transcriptomes using bulk-RNA-seq. All the samples yielded quality reads with the 3 replicates highly correlated and clustered together (Figure 4a). We observed expression of ALT *(gpt, gpt2l)* genes (Supplementary Figure 2) that supported the enzyme colorimetric assays (Table 1), as well as several other important liver metabolism genes in both cell lines. These genes include lipid metabolism genes such as apolipoproteins, *stearoyl-coA desaturase (scdb)* and fatty acid desaturase *(fads2)* and master regulator genes such as *peroxisome proliferator-activated receptor (pparγ)* and *peroxisome proliferator-activated receptor gamma coactivator 1a (pgc1a* or *ppargc1a)* (Figure 4b). The expression of these genes indicates active lipid metabolism in the cell lines (16). To broadly study the categories of genes that are differentially regulated between the two cell lines, we performed differential gene expression analysis. We observed 2472 downregulated genes and 2791 upregulated genes in SFL as compared to CFL. The genes upregulated in SFL were enriched for GO terms ‘lipid metabolic process’, ‘lipid catabolic process’ and several immune system related GO terms such as ‘complement activation’ and ‘activation of immune response’. For instance, genes under the ‘lipid catabolic process’ such as *lipase εb (lipeb), carnitine palmitoyltransferase 1a (cpt1aa), phospholipase D1 (pld1b)* are all upregulated in SFL as compared to CFL. These results reflect the biology of these fishes in vivo, as it has been shown that surface fish undergo higher lipid breakdown while cavefish synthesize more lipid and surface fish have an increased innate immune response compared to cavefish (5,7,14).

**Figure 4:**
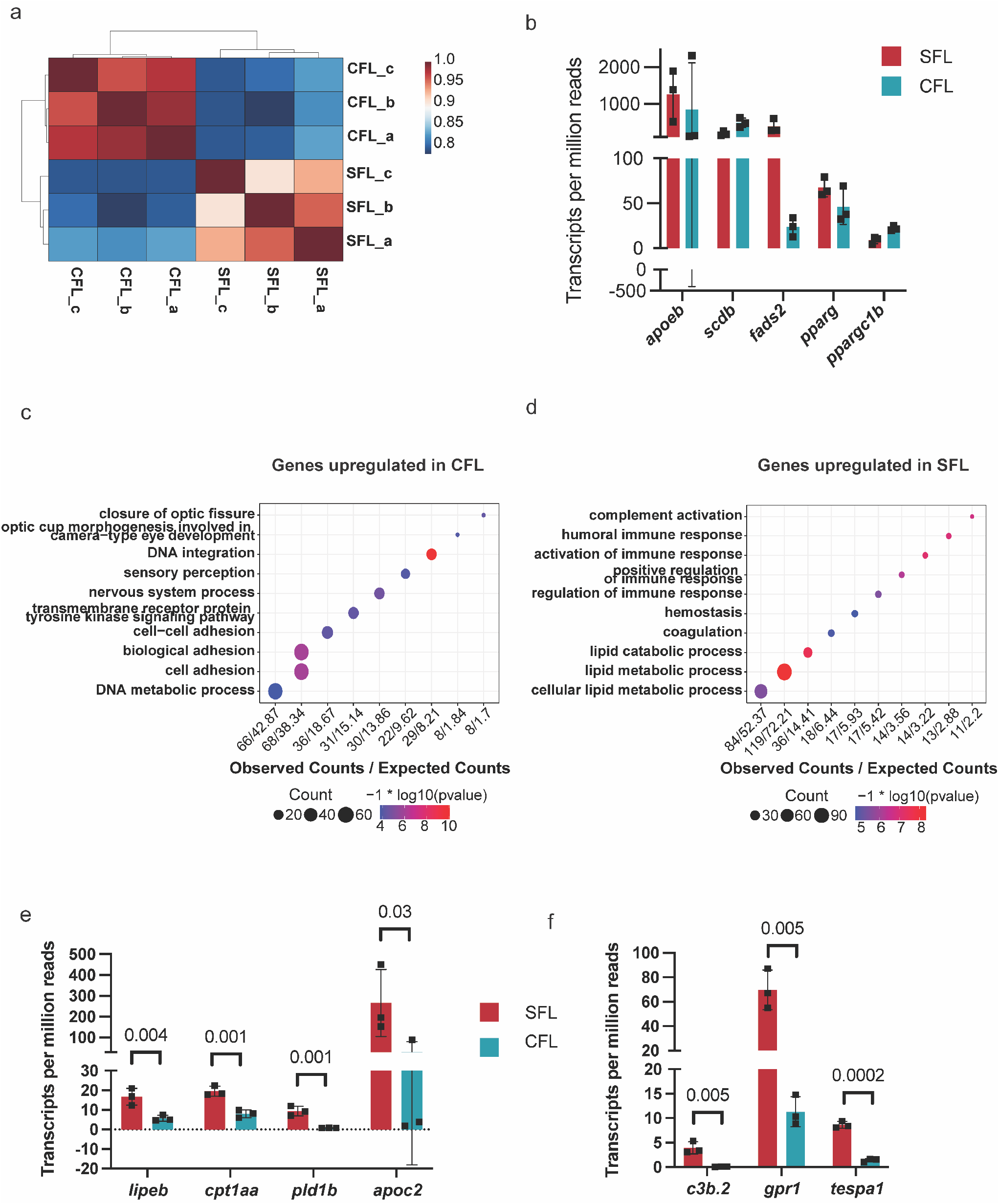
(a) Spearman correlation coefficient heatmap for all samples. (b) Expression levels for some key liver metabolism genes. (c) GO terms enriched among genes upregulated in SFL vs CFL. (d) GO terms enriched among genes downregulated in SFL vs CFL. (e) Genes belonging to ‘lipid catabolic process’ that are upregulated in SFL. (f) Genes belonging to ‘activation of immune response’ that are upregulated in SFL. The bar graphs represent standard deviation around mean values.

### SFL and CFL can be used for generating transgenics

Through the availability of robust gene editing techniques, it has become possible to easily generate gene knockouts in cell lines. Furthermore, cell lines provide an efficient platform for protein overexpression for the study of cell biology and to perform biochemical assays. Thus, to maximize the utility of these cell lines, we tested their ability to be transfected/electroporated. We transfected CFL with a plasmid that expresses GFP under the CMV promoter. We attained good success with transgene expression using electroporation with Lonza nucleofection in CFL (Figure 5). Furthermore, to aid in generation of stable cell lines, we performed kill curves to determine the minimum concentration of G418 needed to kill untransfected cells without affecting the transfected cells. Concentrations ranging from 0μg/ml to 1200μg/ml were tested. We observed complete cell death with 600μg/ml concentration.

**Figure 5:**
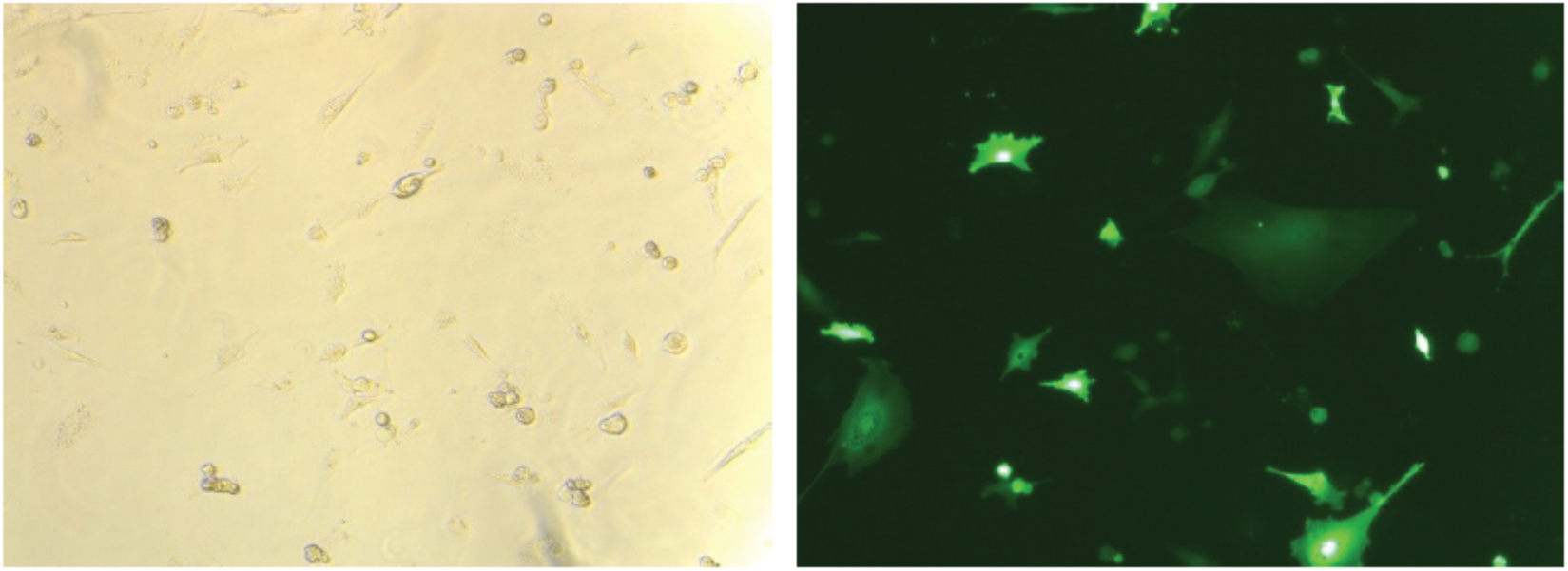
Brightfield and Fluorescence images of SFL electroporated with pMaxGFP plasmid.

These experiments show that the cell lines can be transfected using standard transfection protocols and will increase the utility of these cell lines to study liver metabolism in cavefish.

## Conclusions

In this study we have generated, to our knowledge, the first cell lines from *Astyanax mexicanus.* The liver derived cell lines are adherent and have epithelial/endodermal-like morphology and are easily maintained and sub-cultured in standard tissue culture conditions. The cells express liver enzyme alanine aminotransferase as well as many other key liver metabolism genes. The energy metabolism parameters mirror the in vivo liver physiology of these fish. We also show that these cells are amenable to electroporation that further enhances their utility as in vitro models. The cell lines thus serve as excellent in vitro systems to study and elucidate metabolic features governing the adaptation of cavefish to nutrient poor cave habitats.

## Author contribution

JK and NRO conceived the project, JK, YW, OK, LO, DT and CZ conducted the experiments with support from YW. HH and JK analyzed the transcriptome data. JK and NR wrote the manuscript and all authors read and approved it.

## Supplementary figures

**Supplementary Figure 1:**
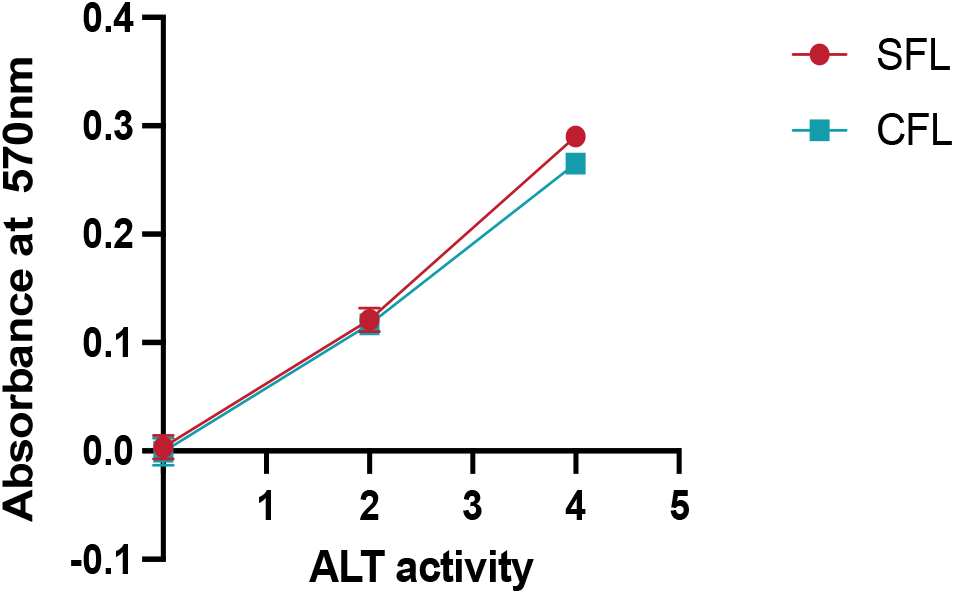
The graphs depict the standard curves for determining ALT enzyme activities using standard colorimetric assay.

**Supplementary figure 2:**
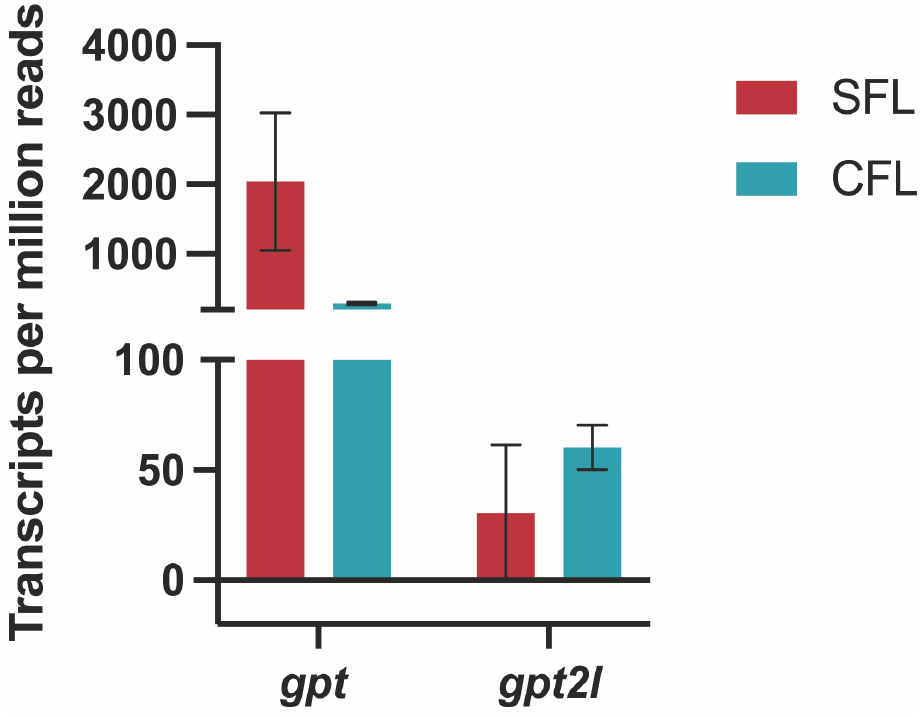
The graph shows expression levels of ALT genes (annotated as Glutamic--Pyruvic Transaminase or *gpt* on Ensembl).

